# MOLECULAR AND QUANTUM MECHANICAL STUDIES OF INTERACTION BETWEEN TETRAHYDROCURCUMIN DERIVATIVE AND PCSK9 Protein TO PROVIDE A BASIS FOR AN ORAL PILL TO REMOVE BAD CHOLESTEROL

**DOI:** 10.1101/2023.07.04.547717

**Authors:** Prakash Vaithyanathan

## Abstract

The interaction between Proprotein convertase subtilisin/kexin type 9 (PCSK9) and low-density lipoprotein receptors responsible for causing atherosclerosis. According to estimates, it causes 60% of fatalities worldwide and is the covert precursor to clinical myocardial infarction (MI), stroke, and CVD. Designing tiny compounds that inhibit PCSK9 from interacting with LDL receptors is the need of the hour. Through bioinformatics-based studies, this study seeks to assess the interactions between a derivative of tetrahydrocurcumin and PCSK9 Protein and compare them to interactions with the literature based studies of standard Atorvastatin. Additionally, comparison research was carried out to examine how the new compound interacts in the active and allosteric regions of PCSK9. The above-mentioned compound, a derivative of Tetrahydrocurcumin, was adjusted and optimized to the level of local minimum energy using the RCSB’s downloaded PDB file 7S5H. By Desmond MD simulation studies, the stability of the non-bonded interactions of the complexes was examined. An affinity of -9.493 kcal/mol for the active site and -8.148 kcal/mol for the allosteric site was observed by docking studies in comparison with the standard molecule, atorvastatin. Also, the MMGBSA value of -50.7142 kcal/mol indicates the Tetrahydrocurcumin derivative binds well compare to the standard, atorvastatin. The Tetrahydro curcumin derivative molecule was able to orient into the active region with the help of Asp238, Thr377, and Ser381 amino acids. In comparison to atorvastatin, the binding affinity was raised by seven H-bonds with six amino acids and one π interaction of Arg295 amino acids of the allosteric site. The Tetrahydro curcumin molecule’s nonbonded interaction was found to be stable for 100 ns by MD simulation tests. This demonstrates that the Tetrahydrocurcumin derivative molecule will prove to be an effective substrate to modify PCSK9 protein behavior.

## Introduction

In the blood, cholesterol has a special role as a crucial precursor for phospholipids, triglycerides, steroids (Bhattarai et al., 2021). Progesterone, estrogen among others are a few of the hormones that are catabolized by cholesterol. When the amount of this essential molecule in the blood exceeds the allowable limits (hyperlipidemia), lipid particles cling to the blood vessel walls and restrict blood flow, which raises the risk of heart attack and stroke. Atherosclerosis is the term used to describe this condition of extremely elevated cholesterol levels (Russo & Jang, 2022; Seidman et al., 2014). In these circumstances, when a human individual has Familial hypercholesterolemia (FH), anti-cholesterol medications are utilized.

Over the past 30 years, statins have typically been the treatment of choice for hyperlipidemia. In the USA, lovastatin (Mevacor), was approved for use by the patients. In subjects who have once experienced a heart attack, statins lower their risk of having another one. In 2020, Basak and Basak Reducing cholesterol, enhancing cardiovascular health, and reducing inflammation are some advantages of using statins. However, according to Ahmad (2014), the main justifications for avoiding statins are their negative effects. Cerivastatin, one of the medications, was taken off the market after it resulted in 52 fatalities. Additionally, rhabdomyolysis, which caused renal system failure, was connected to it. (Pitt and Furberg, 2001).

It was discovered during the quest for a genetically based defect in FH that the majority of cases were attributed to LDL receptor mutations and apolipoprotein genes. In particular, in 2003, scientists discovered that the gene known as PCSK9 has mutations linked to chromosome 1 (Kim, E. J. and Wierzbicki, A. S. 2020). In 90% of people with coronary disease, the PCSK9 protein is overexpressed. According to Schulz R et al. 2021, PCSK9’s catalytic domain, which is comparable to subtilisin, can bind with LDLR EGF-A (epidermal growth factor-like repeats) domain. By inhibiting LDL from adhering to LDLR, Zainab et al. hypothesized that the connection between PCSK9 and LDLR promotes cholesterol buildup in 2021. It is essential to inhibit interaction between PCSK9 and LDLR to prevent CAD (Li et al. 2022; Lin et al., 2018). The two main strategies employed to target PCSK9 rely on small-molecules pharmacotherapies or monoclonal antibodies. (Xu et al. 2019a; Du et al. 2011; Petrilli et al. 2020)

Finding a molecule that binds well with PCSK9 and lower its activity is possible with a profound knowkedge of the molecular structure of PCSK9. Computational methods based on bioinformatics and cheminformatics was employed to find the binding sites of a protein to suppress its activity (2019, Xu et al. Ludington, 2015, and Gioia et al.). The Tetrahydrocurcumin Derivative’s binding profile was studied using dynamic simulation techniques and docking studies in this work.Validation based on realistic models was also carried out to examine the molecules behaviour in the active and allosteric regions.

## Materials and methods

### Software and Modules

To learn about the atomic level interactions and stability of the complex formed between the ligands and proteins, Schrodinger (Zainab R et al. 2021), and Desmond simulation software (Ayaz, P et al. 2023) were used. The molecular mechanics based interaction studies of Tetrahydro Curcumin derivative with PCSK9 protein was carried out using the modules such as protein preparation, Grid generation, Site map prediction, Ligprep, and glide docking (Da Silva et al. 2023 and Fahui Li et al. 2022). The QM-based procedures helped understand the stability of the complex using the modules such as dynamic simulation, simulation quality, and simulation interaction (Bitencourt-Ferreira et al. 2019; Kuki and Nielsen, 2010).

### Protocol for Ligand and Protein preparation

To facilitate the interaction with the PCSK9 protein (7S5H pdb), the tetrahydrocurcumin derivative molecule’s energy has to be minimized to a local energy level ensuring its stability. The preprocessing steps for the protein was carried out by enabling Assign bond order using the CCD database along with the hydrogens substituted (Lima et al. 2022). The PDB file 7S5H was used as a source entry in the preparation routine. Based on the zero-order, the metal and disulfide connections are formed. The pH states generated by the Epik module are 7.0 +/- 2.0. Using the PROPKA optimization approach and sample water orientation, this preprocessed protein was assigned with an optimized H-Bond. The protein was finally minimized using the OPLS4 force field calculation and the heavy atom RMSD coverage was set as 0.30 A. Beyond a distance of 5 A from the PDB ligand, the water molecules present were removed. The LigPrep protocol was used to prepare the tetrahydrocurcumin derivative molecule ready for docking. The Epik algorithm was used to desalt the system by enabling the OPLS4 force field parameters. For the ligand molecule, 32 stereoisomers were produced in total. (Madhavi Sastry et al. 2013)

### Binding pocket analysis using the Grid generation method

The peptide location in the pdb protein 7S5H was taken into account by the receptor grid-generating process as a binding pocket for the tetrahydrocurcumin derivative and atorvastatin ligands. The VdW radius scaling factor is set to 1.0 with 0.25 as the cutoff partial charge. The enclosure box was sized to fit the workspace ligand’s centroid, and the length of the dock ligands was set at 12. Using the site maps procedure with a normal grid size of 4 crop at the regions closer to the allosteric site of 7s5h, the actual Allosteric site was found. An allosteric grid is constructed for docking the molecules based on the site map locations. (2013) (Wang et al.)

### Ligand docking

For the ligand docking, the scaling factor technique was adjusted to 0.80 and the partial charge cutoff (van der Waals radii) to 0.15. The flexible ligand sampling setup was employed along with the imported grid using the extra precision approach (Friesner et al., 2004). All preset functional groups were employed for bias sampling of torsions, along with sample nitrogen inversion, ring conformations, and sample nitrogen inversion. In the end, the docking score was increased by Epik state penalties. For the creation of conformers, the ring sampling energy window was set as 2.5 kcal/mol, the dielectric constant value being a function of distance is set as 2.0 for the purpose of minimization(2002) (Malla et al). To explore the binding energies with the PCSK9 (7S5H) protein, the Prime-MMGBSA technique was also carried out for the better poses of tetrahydrocurcumin molecule and Atorvastatin. According to L. B. Silva et al. (2023), the solvated model and force field protocols were set to VSGB and OPLS4, respectively.

### Dynamic Simulation Stability Protocol

Using the system builder protocol in Desmond software, the stability of the best-docked posture of the tetrahydrocurcumin derivative and Atorvastatin was investigated. The complex tetrahydrocurcumin derivative – 7s5h was solvated firstly using the solvent model of SPC, with orthorhombic boxes as the boundary conditions, and the size of the buffer box was calculated at a distance of 10.0 A. Additionally, Na+ and (Cl-) ions was used to neutralize the complex system and the final volume was minimized based on the complex’s surface occupation. The protocol for molecular dynamic simulation was applied to the solvated model. A chosen energy gap value of 1.2 was used with a fixed simulation time of 100 ns and a 100 ps interval for the trajectory recording. A total of 1000 frames were produced in an NPT ensemble class, with the model relaxed before to simulation, at a a pressure of 1.01325 bar and at a temperature of 300 k.

The finished complex simulation was loaded, and the average block length for the analysis of the trajectory quality was 10.0 ps. Using the parameters for Potential energy (kcal/mol), Total energy (kcal/mol), pressure (bar), temperature (k), and volume (A3), graphs can be created using the simulation quality analysis technique. The simulation interaction diagram module finally determined a complex’s RMSD, RMSF, and Torsion values with a time scale set as 100 ns.

## Result and Discussion

### Grid Box Optimization

The energy values of protein 7S5H and the ligand molecules were optimized to a local minima using the preparation techniques to make it stable for the docking investigations. The overlaid picture of a protein before and after minimization is shown in Fig 1. The binding area loop amino acids comprised of ASP192, ASP238, and GLY 384 were located close to where the most significant conformational alterations were seen. Similarly to this, the original state and the energy-minimized state of the Tetrahydrocurcumin Derivative molecule (Pincock & Torupka, 1969) were superimposed. The cyclic ring fragments with the RMSD of 0.4570 showed configurational alterations. The ultimate energy for the protein was found to be –2272.338 kcal/mol and for the molecule it was -86.852 kcal/mol.

**Fig 1.**
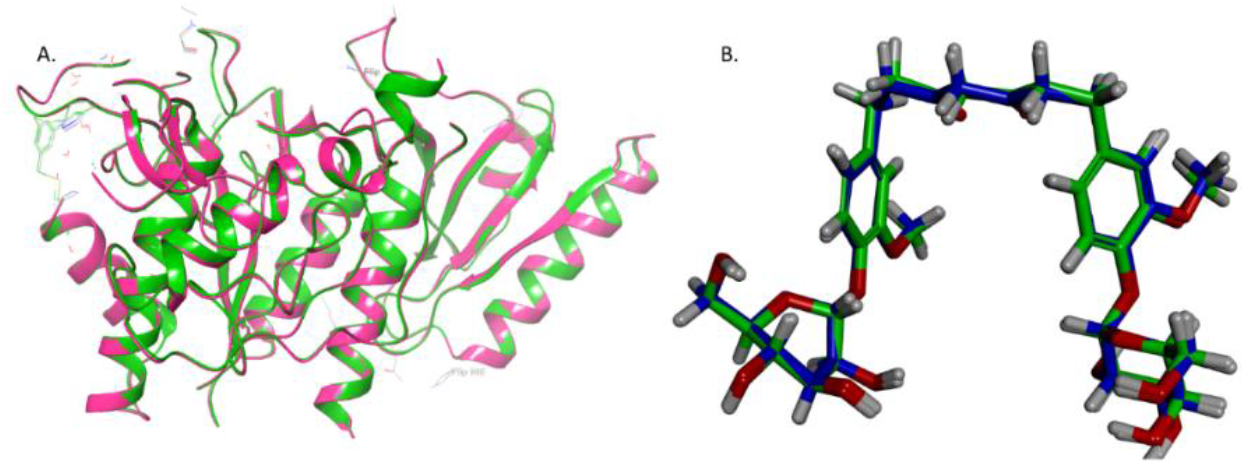
Secondary structure of protein 7S5H before (green) and after (Pink) siperimposed after minimization. Superimposed tetrahydrocurcumin molecule derivative before (green) and after (red) minimization.

The binding grid pockets, active site (with X, Y, Z coordinates set as -17.16, 19.08, and - 6.2)and the allosteric site (X, Y, Z coordinates set as -22.5, -20.67, and 11.3) and 10 Å is set as the radius in both the sites) (Fig 2A) to bind the tetrahydrocurcumin derivative and atorvastatin.

**Fig 2.**
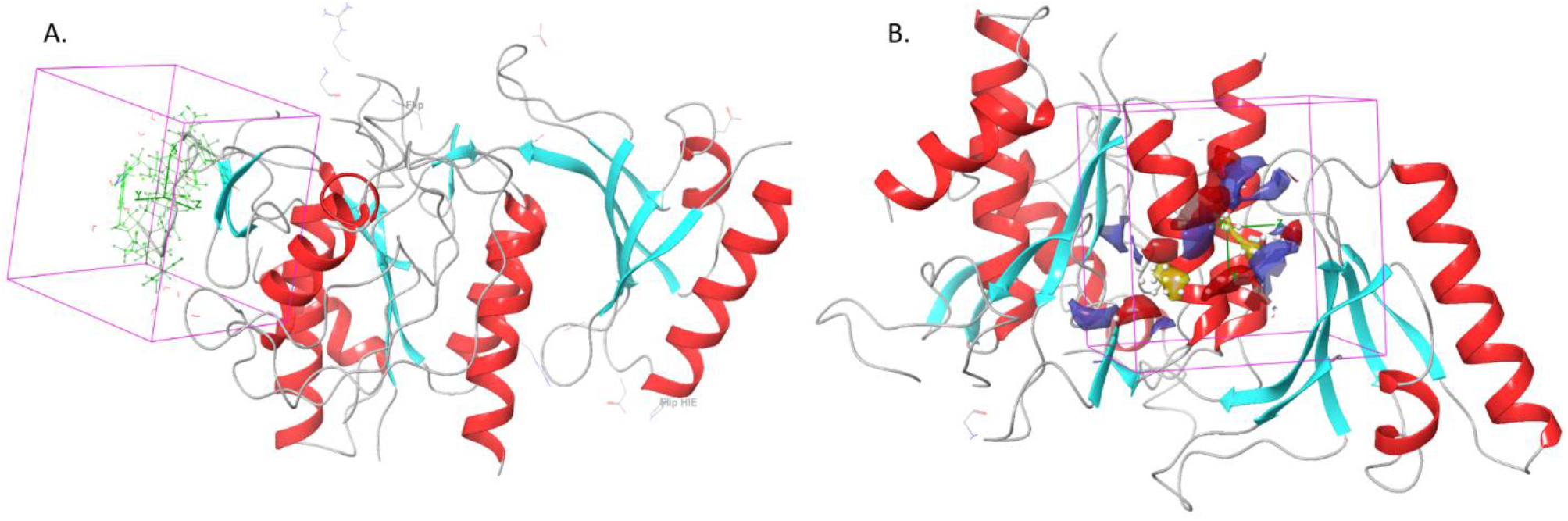
**A**. 7S5H protein’s Active site. B. 7S5H protein’s Allosteric site

**Fig 3.**
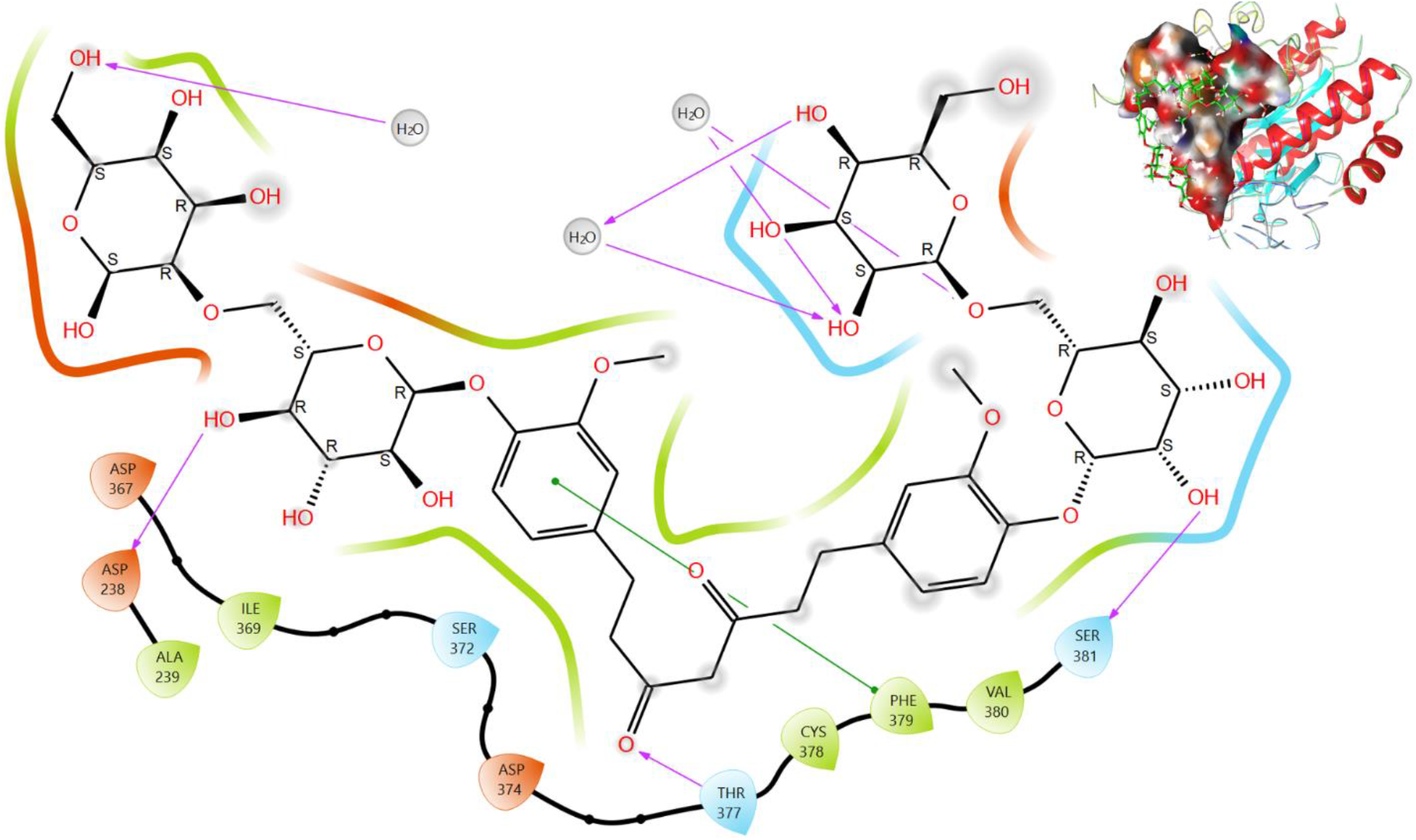
2D representation of the interacting amino acids with the atoms of tetrahydrocurcumin derivative in the PDB active site of 7S5H protein

**Fig 4.**
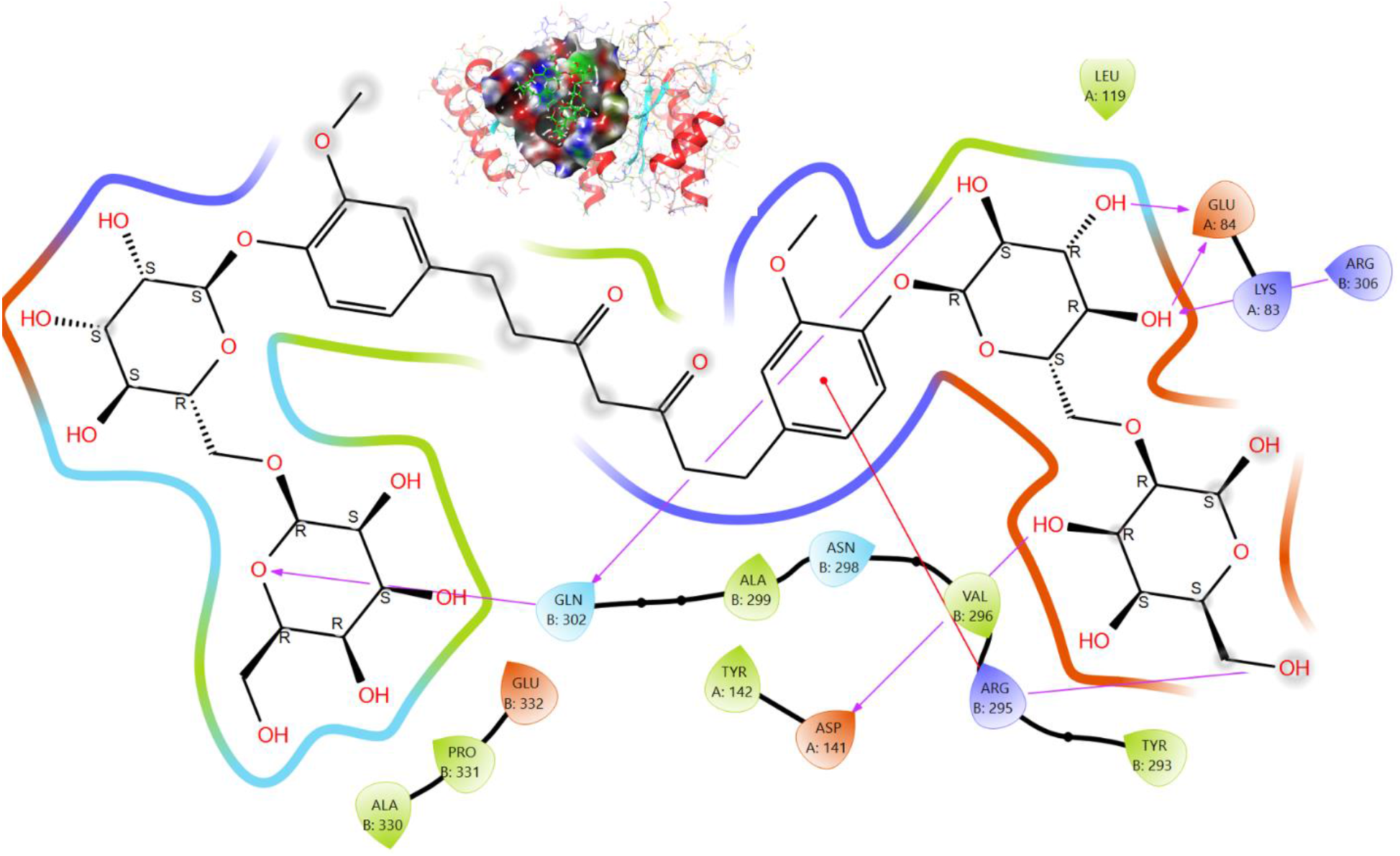
2D interaction of amino acids of pdb allosteric site of 7S5H protein with the atoms of tetrahydrocurcumin derivative.

### Tetrahydrocurcumin derivative and atorvastatin docking analysis

Glide docking procedures demonstrated the formation of good poses of atorvastatin and tetrahydrocurcumin derivative in the interiors of the binding regions. The energy of the tetrahydrocurcumin derivative molecule in both sites was -9.493 kcal/mol and -8.148 kcal/mol and were almost twice that of atorvastatin. The tetrahydrocurcumin derivative molecule was supported well by the water molecules to help bind better when compared with atorvastatin and the hydroxyl groups of the tetrahydrocurcumin derivative help to generate 5 H-Bonds with 3 water molecules and four H-bonds with Asp238, Thr377, and Ser381 amino acids and with the - OH group that constitutes the sugars. In addition, 3 polar residues Thr377, Ser372, Ser381, 5 amino acids Ala239, Ile369, Cys378, Val380, Phe379 (Hydrophobic), 3 negatively charged amino acids Asp238, Asp367, and Asp374 help modify the configuration of the tetrahydrocurcumin derivative for good binding. In addition, pi-pi stacking interactions with Phe379 and a benzene ring of the core skeleton of the tetrahydrocurcumin derivative molecule as well. Further, the binding of tetrahydrocurcumin derivative was confirmed by Mmgbsa with an active site binding energy level of -50.71 kcal/mol and an allosteric binding energy level of - 37.48 kcal/ mol.

**Table 1.**
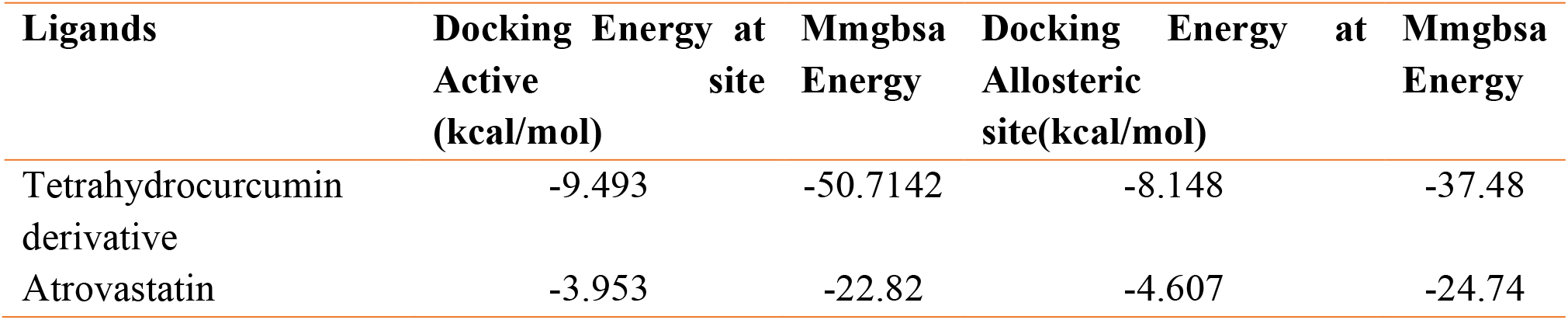
Dock energy values of Tetrahydrocurcumin derivative and atorvastatin molecules

| Ligands | Docking Energy at Active site (kcal/mol) | Mmgbsa Energy | Docking Energy at Allosteric site(kcal/mol) | Mmgbsa Energy |
| --- | --- | --- | --- | --- |
| Tetrahydrocurcumin derivative | -9.493 | -50.7142 | -8.148 | -37.48 |
| Atrovastatin | -3.953 | -22.82 | -4.607 | -24.74 |

The residues at the allosteric site also generated multiple links. Seven amino acids of 7S5H such as Glu A: 84, Arg B:306, Arg B: 295, Gln B:302, and AspA:141 helped to form H-bonds which turned the tetrahydrocurcumin derivative to make proper confirmation and interact hydrophobically with other amino acids (Leu A:119, Tyr B:293, Val B:296, Ala B:299, TyrA:142, Pro B:331, and Ala B330), Positively charged residue (ArgB295, Lys A83, Arg B306), Polar (Gln B:302, and Asn B:298) and Negatively charged (Glu B:332 and Asp A: 141) interactions.

The standard drug atorvastatin’s interaction in the active and allosteric site of the protein is shown in Fig 5. The image showed that the common atorvastatin created four different sorts of interactions, with the hydrogen bond creation specifically involving the carboxylic group and the aliphatic chain --OH group. The aniline ring orientation is supported by a contact between the negatively charged amino acid Asp238 and the molecule’s R-conFigd -OH group, that interacts with the amino acid Ser381. The pyrrole ring’s location in the active site is aided by the amino acid Phe379’s -interaction with the molecule’s benzene ring system.

**Fig 5.**
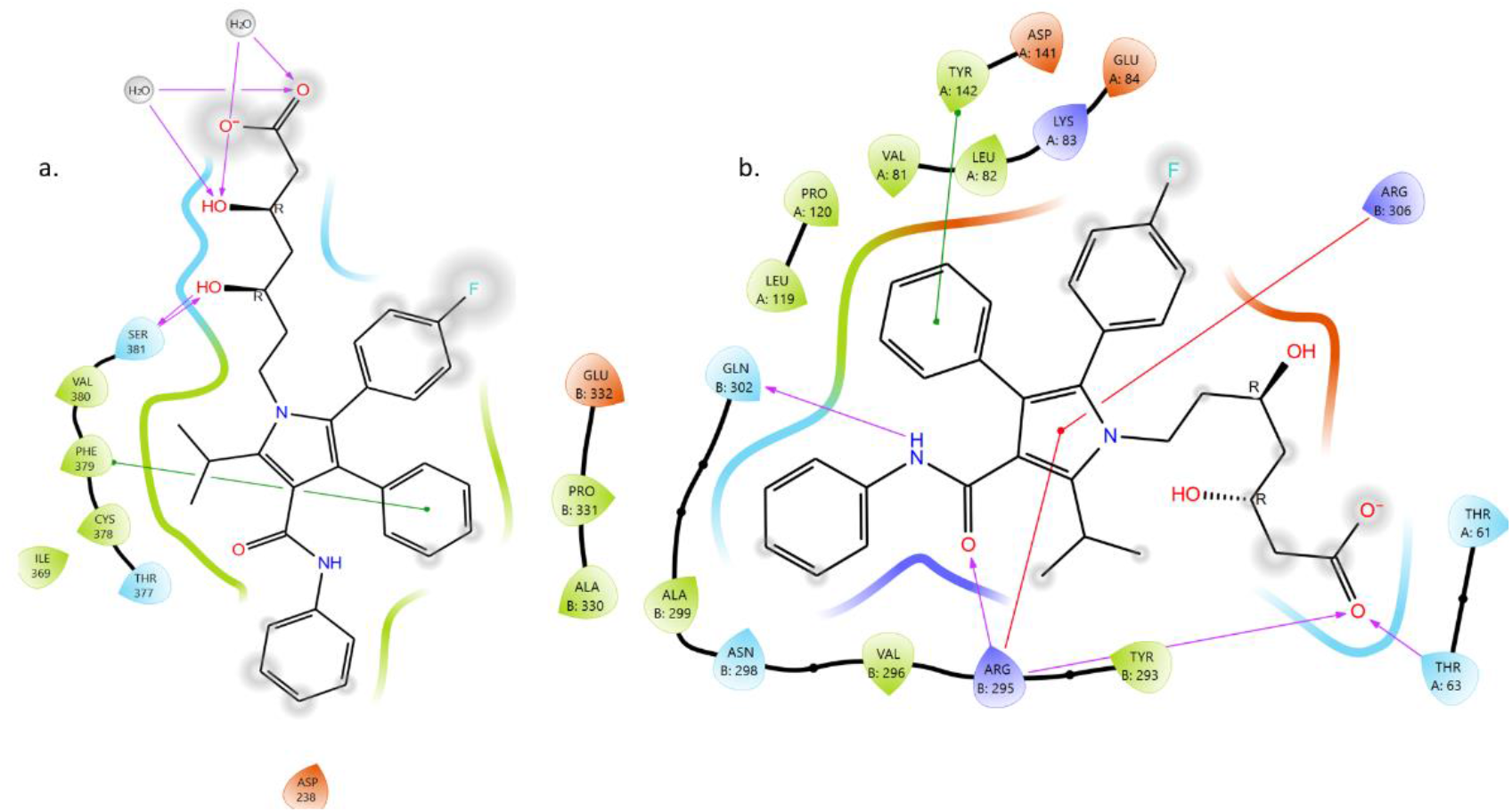
a. Atorvastatin’s binding with the protein’s active site. b. Atorvastatin’s binding with the protein’s allosteric site

The H-bond and π-π interactions were proven to be crucial due to the allosteric binding of atorvastatin, as illustrated in Fig. 5b. The fixing of the aniline ring is supported by the hydrophobic amino acids AlaB299, the polar amino acids AsnB298 and Gln302. The benzene ring position is assisted by LeuA82, ValA81, ProA120, and LeuA119, and the core centre pyrrole moiety is fixed to orient inside the allosteric site by the -cationic formation with ArgB295 and ArgB306 (Fig. 5b) (Irfan N et al., 2023).

### Analysis of stability of Dynamic simulation

It is not possible to establish the potential efficacy and bond formation of the Tetrahydro Curcumin derivative just by interaction studies. Using simulation studies from MD, it should be examined for dynamic stability. Galvo Lopes and coworkers, 2023; Navabshan and coworkers, 2021). Water was used to dissolve the Tetrahydro Curcumin derivative and Atorvastatin molecule complexes, and Na+ and Cl-ions were used to neutralise them. For the Tetrahydrocurcumin derivative active site binding complex, the system builder protocol included 12695 water molecules, whereas 12350 water molecules were added for the Atorvastatin protein complex system for 100ns simulation experiments (Fig. 6).The additional 345 water molecules were added in the complex of 7S5H protein compared to the standard atorvastatin drug complex and the addition was made to cover the extra volume generated by the extended substitution of Tetrahydro Curcumin derivative.

**Fig 6.**
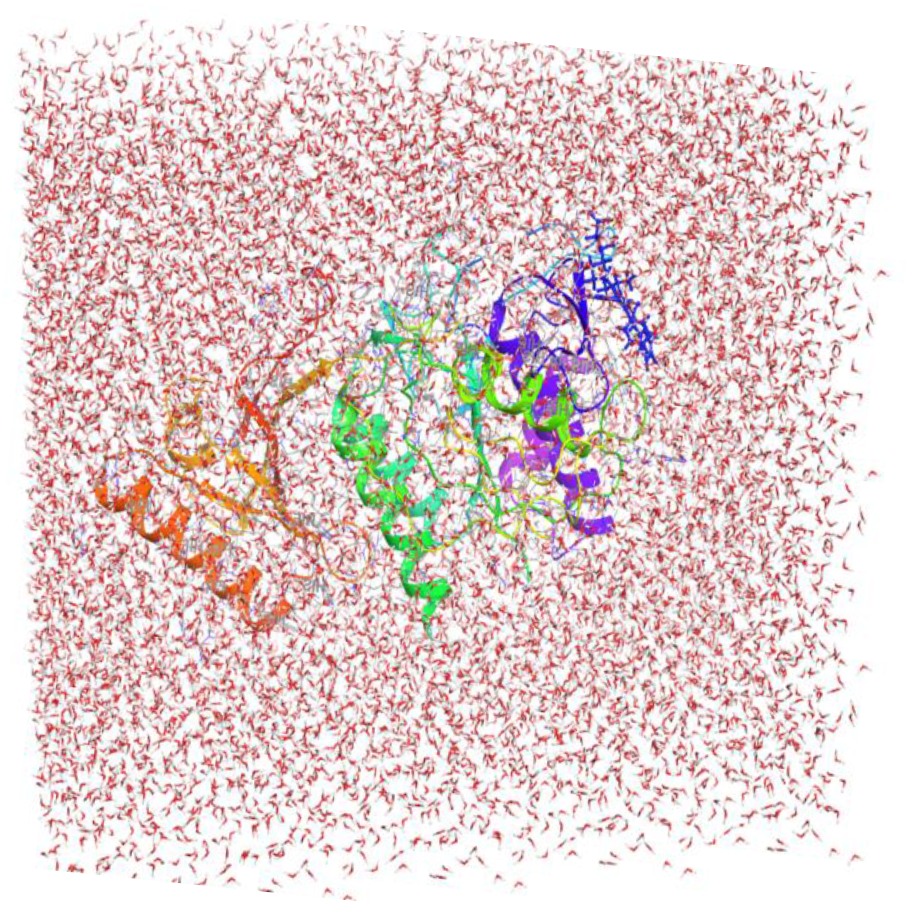
Tetrahydrocurcumin derivative binding to the active site in a solvated model.

### Analysis of the system’s quality and a dynamic interaction diagram

The quality of the molecular dynamic simulation run was examined, and the parameter values were found to be within a reasonable range. The complex simulation quality, as shown in Fig. 7, has reliable values for volume (349810.091 ± 361.968), pressure (1.08361 bar), temperature (298.680 ± 0.620), and potential energy (117704.585 ± 78.559). The total energy values of -96208.168 kcal/mol showed that the 7S5H protein-tetrahydrocurcumin combination remained stable during the course of the simulation’s 100 ns.

**Fig 7.**
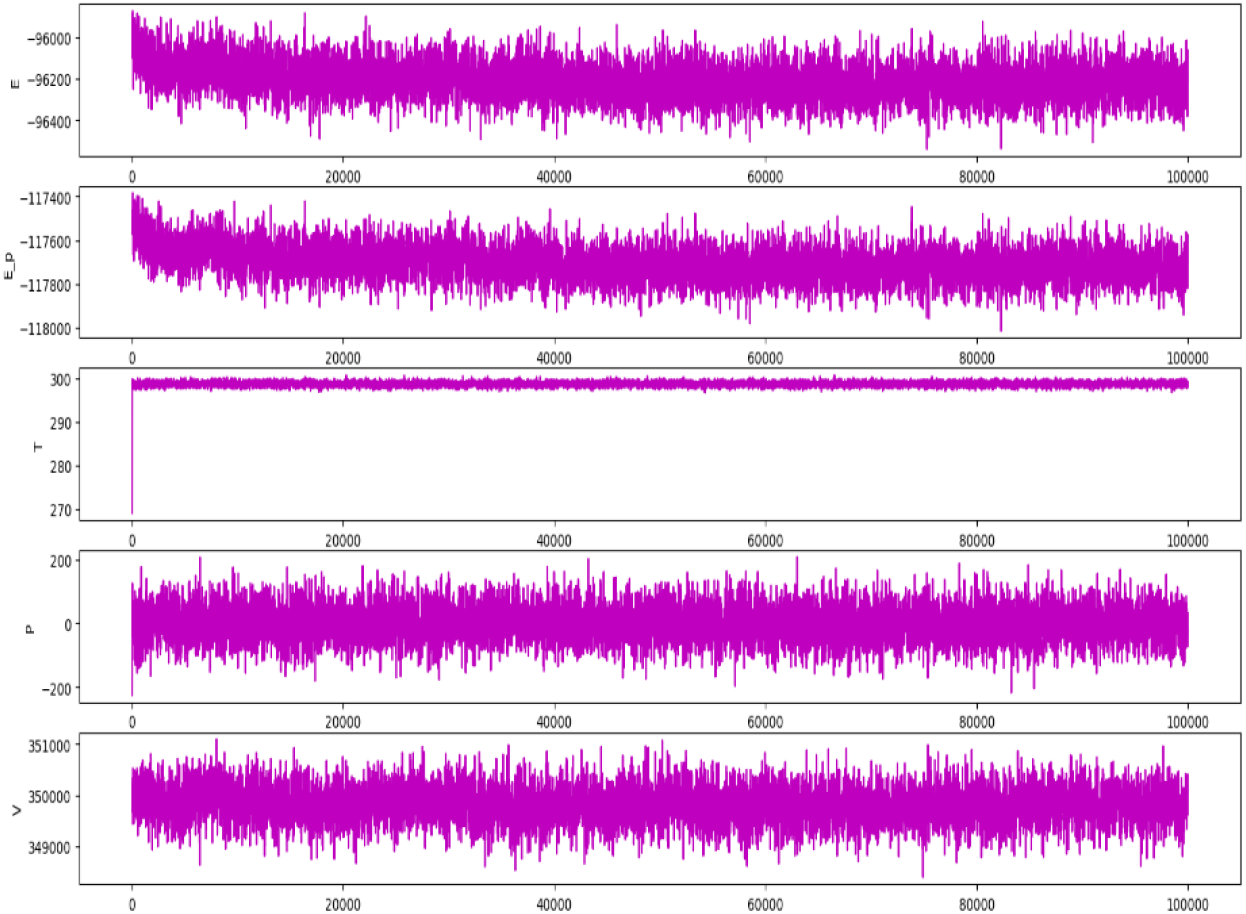
Tetrahydro curcumin-protein complex simulation run quality analysis report for 100 ns

### Root Mean Squared Deviation RMSD, Root Mean Squared Fluctuation RMSF, and Protein-ligand Contacts in the active site

The 7S5H-tetrahydrocurcumin derivative complex began to stabilise at 10 nanoseconds, with an RMSD range of 1.50 Å - 1.80 Å (Fig. 8a), according to the RMSD graph. The RMSD values fluctuated little between the intervals of 10 and 34 ns, then abruptly increased between 34 and 38 ns. The complex was observed to be fully stabilised based on the RMSD measured after the 39th ns. Atorvastatin-7S5H complex RMSD was displayed in Fig. 8b, demonstrating that the complex was unstable beyond 80 ns.

**Fig 8.**
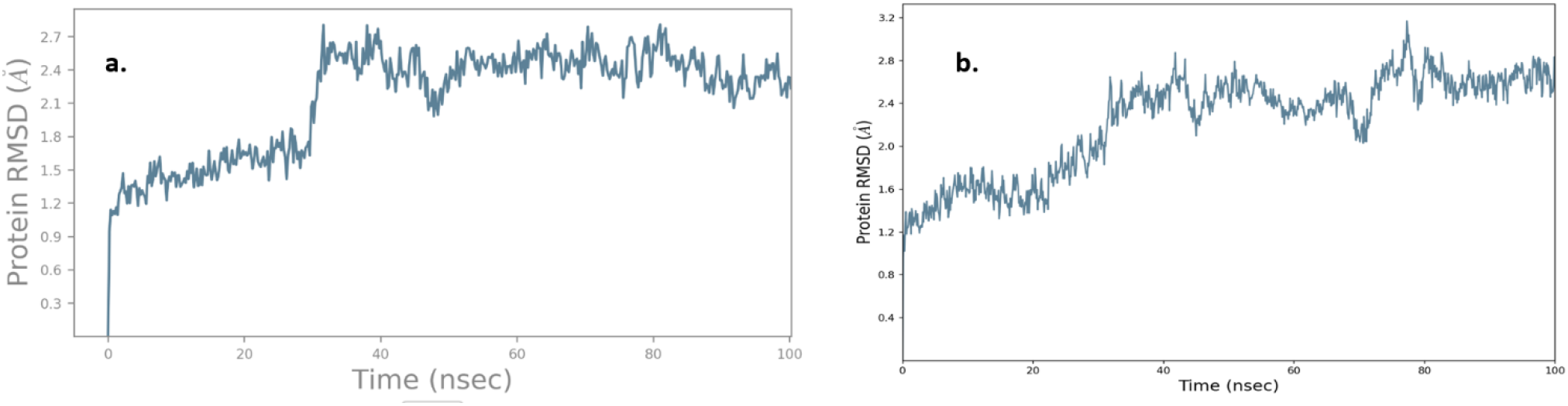
a. RMSD of different conformations of 7S5H-Tetrahydrocurcummine tetracyclo at 100ns. b. RMSD of different conformation of 7S5H-Atrovastatin at 100ns.

During the simulation run, the interactions of the 7S5H-tetra hydro curcumin complex, shown in Fig. 9, caused the molecule to remain stable over the 100ns time duration. During the simulation, the three hydroxyl groups of the glucose moiety and the four water molecules experienced substantial contact due to the negatively charged Asp B:238. The two oxygen atoms of the Tetrahydro Curcumin derivative bonded with the polar amino acid His B:391 by the help of a water molecule. The stiff ring is positioned inside the active binding pocket of the 7S5H protein thanks to the direct π-π hydrophobic stacking contact generated by the B: 379. The complex is stabilised and its oscillations are lessened by these interactions and configurations of amino acids.

**Fig 9.**
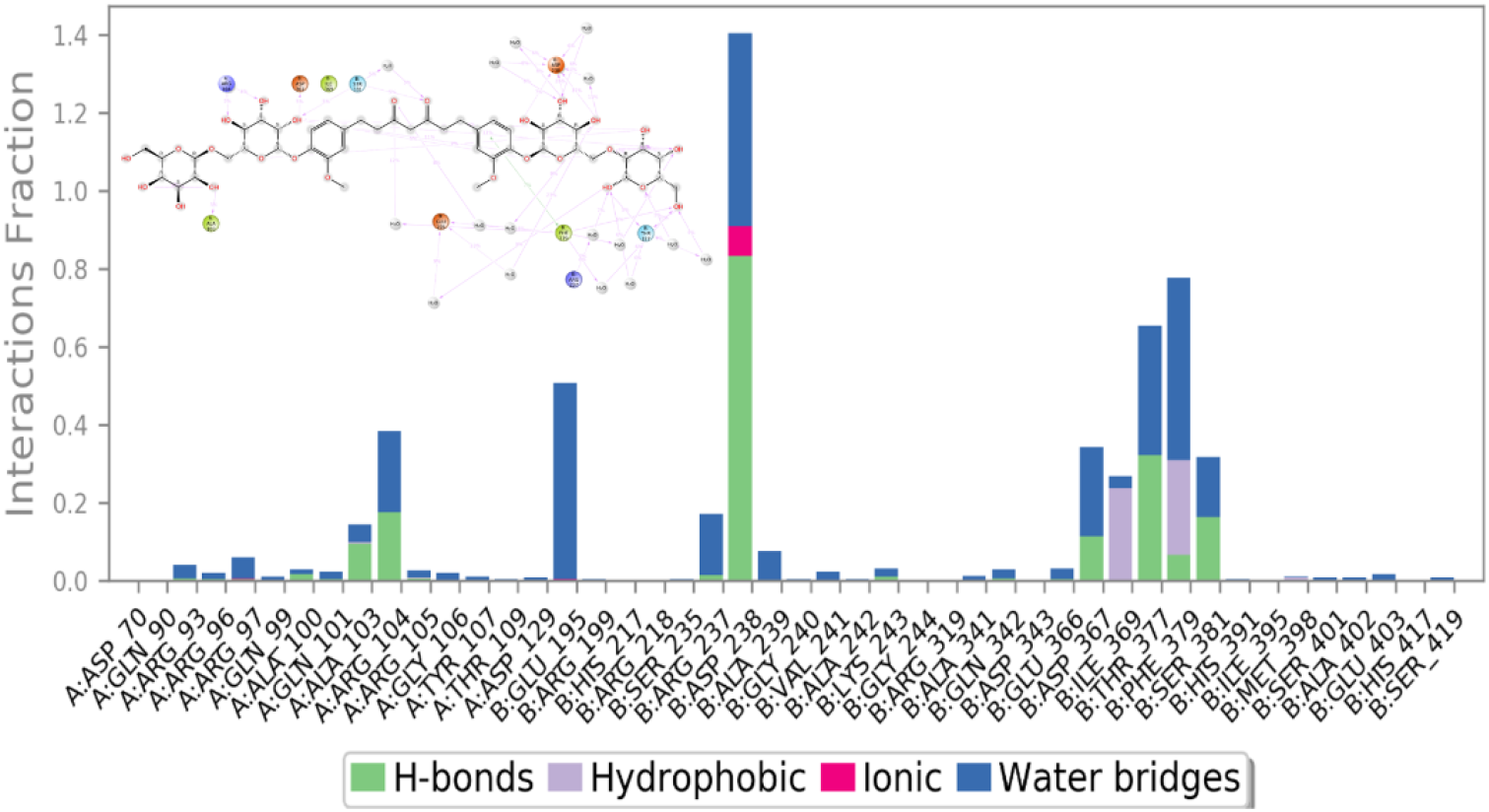
Interaction map of the Tetrahydrocurcumin derivative molecule during the 100ns simulation

As shown in Fig. 10, the atorvastatin-protein complex exhibits more RMS fluctuation of amino acids of 7s5h than does a derivative of tetrahydrocurcumin. Particularly, the amino acids 132, 143, and 250 varied by more than 5 Å, which caused atorvastatin to be less stable with 7S5H protein. Tetrahydrocurcumin derivative atoms established many bonds with residues 10, 35–49, 72, 108, 125–127, 246-254, and 247–312 that resulted in the stable interaction with the 7S5H protein.

**Fig 10.**
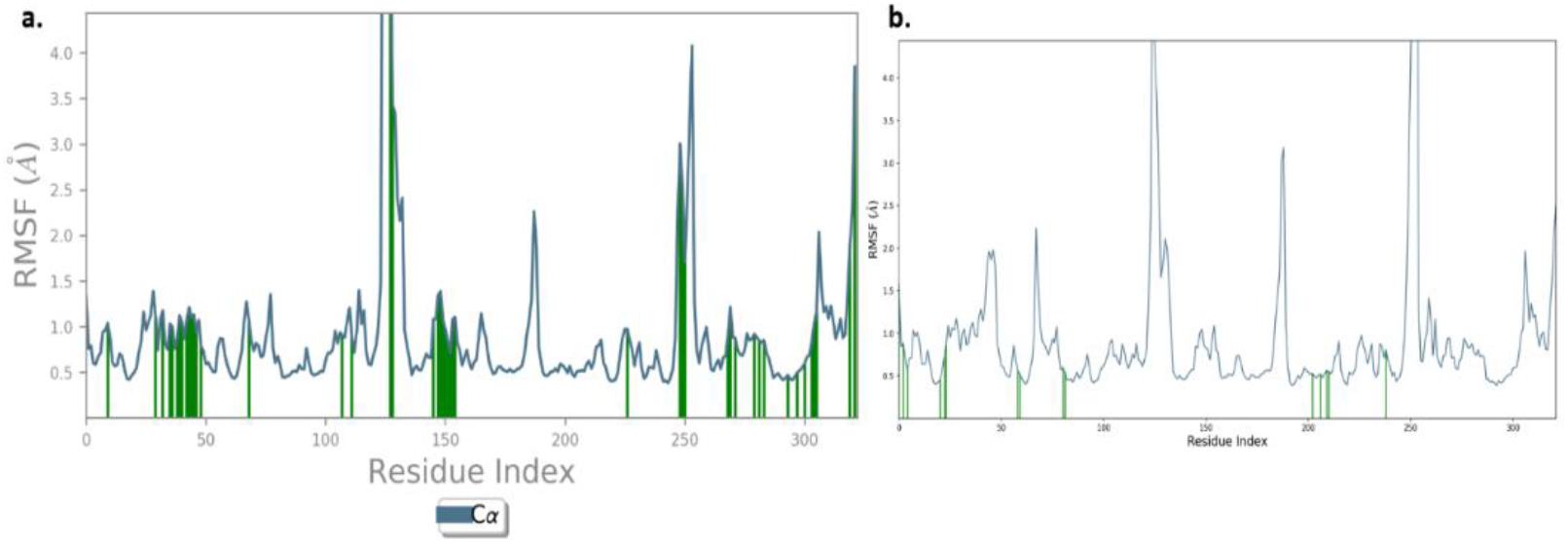
a. Tetrahydrocurcumin derivative-7S5H protein’s RMSF. b. Atorvastatin’s RMSF

### Root Mean Squared Deviation RMSD, Root Mean Squared Fluctuation RMSF, and Protein-ligand Contacts in the allosteric site

Figure 11 analyse and depict the stability of the Tetrahydrocurcumin derivative and Atorvastatin. The 7S5H-tetra hydro curcumin complex began to stabilise from the eighth nanosecond forward, followed by a fluctuation in the fourteenth nanosecond, and then became stable (Fig. 11a), with the deviation range being 1.50 - 2.0 Å. This is shown by the RMSD of allosterically binding molecules. With a variance of 2.0-3.2 Å ns, the atorvastatin (Fig. 11b) molecule was not getting stabilised up to 71 ns.

**Fig 11.**
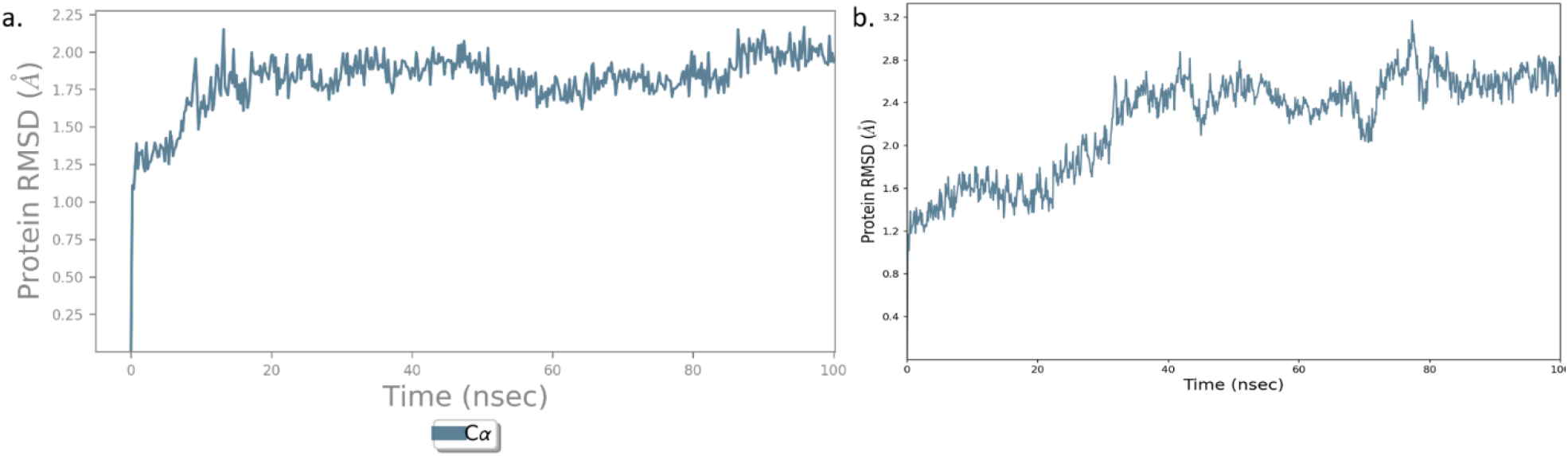
a. RMSD of multiple 7S5H-Tetrahydro Curcumin conformations at 100ns inside the allosteric location. b. RMSD of 7S5H-Atorvastatin conformations at 100ns inside the allosteric site.

During the simulation, 33 residues, 13 from the A chain and 20 from the B chain, were implicated in the allosteric binding of the 7S5H-Tetrahydrocurcumin derivative, which helped decrease the conformational fluctuations of the side chain hydrogen and keep the complex stable (Fig 12a). Specifically, the B chain residues Tyr 293 and Arg 357 play an important role in the protein’s folding stability. Only a few amino acids were implicated in protein stabilisation in the atorvastatin binding (Fig 12b). The allosteric site of Tetrahydro Curcumin derivative and atorvastatin complexes was studied for residue fluctuation. The green colour line represents the residues that establish bonds with the fragment of molecules. The fluctuation of residues caused by Tetrahydrocurcumin derivative fragment binding is depicted in Fig 13a. Multiple contacts were created by the amino acid residues 2 to 11, 21 to 24, 81-84, and 199 to 260 to help hold the residues together and form a suitable stable secondary structure of the complex. Only the pyran fragment in Atorvastatin made a few contacts with the allosteric site residues.

**Fig 12.**
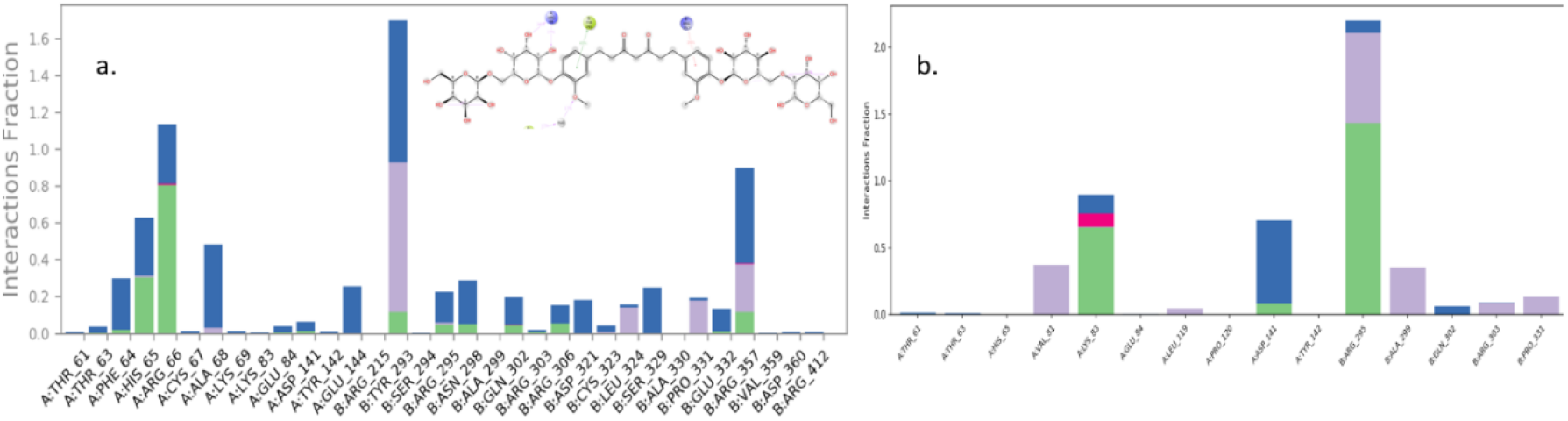
a. Allosteric site residue interaction developed during the 100 nanoseconds simulation of 7S5H-Tetrahydrocurcumin derivative. b. Allosteric site residue interaction developed during the 100 nanoseconds simulation of 7S5H-Atrovastatin.

**Fig 13.**
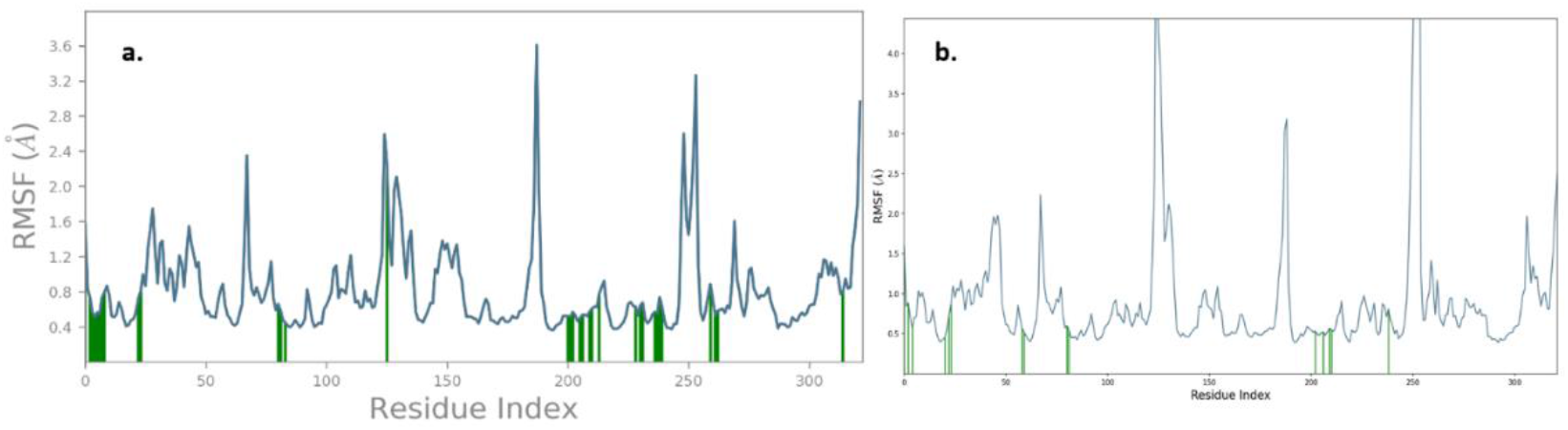
a. RMSF of Allosteric site residue interaction developed during the 100 nanoseconds simulation of 7S5H-Tetrahydrocurcumin derivative. b. RMSF of Allosteric site residue interaction developed during the 100 nanoseconds simulation of 7S5H-Atrovastatin.

When compared to normal atorvastatin, Tetrahydrocurcumin derivative bind effectively in the active site and allosteric site. The MD studies validated the 7S5H protein-tetrahydro curcumin complex’s stability while binding. This study also shows that Tetrahydrocurcumin derivative can be a potential molecule for inhibiting the PCSK9 protein (7S5H) and assisting in the development of an oral pill to remove bad cholesterol.

## Conclusion

The results of this work’s interaction analysis and stability investigations suggested that the molecule Tetrahydrocurcumin derivative was better positioned to be a lead molecule to develop an oral pill to remove bad cholesterol. Specifically, substituting OH groups on the tetrahydropyran ring improves affinity to active and allosteric site amino acids. The MD stability analysis supports the establishment of unbreakable non-bonded connections that keep the molecule stable within the protein. The fabrication of the Tetrahydrocurcumin derivative’s stiff skeleton may boost activity to aid in the treatment of cardiovascular-related issues.

## Conflict of Interest

There is no conflict of Interest

## Notes

### Competing Interest Statement

The authors have declared no competing interest.

